# Adapting Genotyping-by-Sequencing for Rice F2 Populations

**DOI:** 10.1101/055798

**Authors:** Tomoyuki Furuta, Motoyuki Ashikari, Kshirod K. Jena, Kazuyuki Doi, Stefan Reuscher

**Affiliations:** Nagoya University, Bioscience and Biotechnology Center, 464-8601 Nagoya, JAPAN; International Rice Research Institute, DAPO Box 7777, Manila, PHILIPPINES; Nagoya University, Associated Field Science and Research Center Togo Field, 470-0151 Nagoya JAPAN

**Keywords:** Genotyping-by-sequencing, SNP marker, rice breeding, trait mapping

## Abstract

Rapid and cost-effective genotyping of large mapping populations can be achieved by sequencing a reduced representation of the genome of every individual in a given population and using that information to generate genetic markers. A customized genotyping-by-sequencing (GBS) pipeline was developed to genotype a rice F2 population from a cross of *Oryza sativa* ssp. *japonica* cv. Nipponbare and the African wild rice species *Oryza longistaminata*. While most GBS pipelines aim to analyze mainly homozygous populations we attempted to genotype a highly heterozygous F2 population. We show how species-and population-specific improvements of established protocols can drastically increase sample throughput and genotype quality. Using as few as 50,000 reads for some individuals (134,000 reads on average) we were able to generate up to 8,154 informative SNP markers in 1,081 F2 individuals. Additionally, the effects of enzyme choice, read coverage and data post-processing are evaluated. Using GBS-derived markers we were able to assemble a genetic map of 1,536 cM. To demonstrate the usefulness of our GBS pipeline we determined QTL for the number of tillers. We were able to map four QTLs to chromosomes 1, 3, 4 and 8 and confirm their effects using introgression lines. We provide an example of how to successfully use GBS with heterozygous F2 populations. By using the comparatively low-cost MiSeq platform we show that the GBS method is flexible and cost-effective even for smaller laboratories

## INTRODUCTION

The advances in sequencing technology have drastically improved our ability to determine and simultaneously genotype genetic markers (Davey et al. 2011). The enormous number of short (50 to 200 bp) reads produced by sequencing platforms has drastically reduced the costs and time associated with DNA sequencing. Those advances may be utilized in whole-genome resequencing approaches to generate a collection of reads from untargeted sites in the genome (Takagi et al. 2013; Duitama et al. 2015). Other approaches aim at reducing the complexity of the genome by sequencing only a targeted fraction of the genome. Those genotyping-by-sequencing (GBS) approaches were successful in generating tens of thousands of markers even in plant species with large and repetitive genomes, like maize, wheat or barley (Poland et al. 2012; Romay et al. 2013) or in more heterozygous animal species like cattle or pig (Gualdrón Duarte et al. 2013; Donato et al. 2013).

It was shown that GBS can be used as a fast and cost-effective tool in population genetics, QTL (quantitative trait locus) discovery, high-resolution mapping and genomic selection (Spindel et al. 2013; Huang et al. 2014; Rabbi et al. 2014; Elmer et al. 2015; Lin et al. 2015; Burrell et al. 2015; Begum et al. 2015). Since GBS data typically generates relatively dense marker data a popular analysis choice is a genome-wide association study (GWAS) (He et al. 2014; Begum et al. 2015; Sonah et al. 2015). This kind of study employs a panel of cultivars or varieties. In addition, there are some examples of QTL analyses using bi-parental populations combined with GBS (Spindel et al. 2013; Honsdorf et al. 2014). In those studies recombinant inbreed lines that already underwent several rounds of selfing were used to detect QTLs. There are also examples of the use of GBS to genotype less fixed populations, like F2s (Pootakham et al. 2015; Rowan et al. 2015). In many cases desirable traits are found only in wild relatives or are spread across diverse elite cultivars. The application of GBS to genotype F2s or breeding materials will greatly facilitate gene discovery and marker-assisted selection in breeding projects.

While GBS certainly has huge benefits for scientists and the breeding community, there are some inherent drawbacks to which no universal solution has been found yet (Poland and Rife 2012; He et al. 2014). The data produced by GBS and similar strategies has many missing datapoints compared to datasets from classical, “manually” produced genetic marker data or chip-based systems. Furthermore, there is a considerable error-rate associated with GBS-derived genotypes. Both of these issues can be dealt with at the cost of intensive post-processing, data correction and imputation, which is time consuming and requires specific bioinformatics attention. Also, for each GBS project the researcher has to balance the cost of the sequencing platform with the goal of generating high enough read coverage and in turn marker resolution for the intended analysis. Most GBS strategies aim to sequence only a defined fraction of the whole genome to reduce the number of reads necessary for adequate per-marker read coverage. A common approach is the use of one or two REs to produce fragments with defined endpoints instead of random shearing of input DNA. A recent protocol (Elshire et al. 2011) uses a combination of a restriction enzyme (RE) with a 6 bp recognition sequence to target specific sites in the genome and a RE with a more common 4 bp recognition sequence to generate fragments of suitable length. It was also shown that the choice of RE can influence sequencing results (Heffelfinger et al. 2014). Another common strategy to reduce sequencing costs is the use of multiplexed libraries. By ligating a sample-specific, unique adapter sequence (also called a barcode) to the DNA fragments before pooling and library preparation DNA from multiple individuals may be processed in a single library. Currently, between 96-fold and 384-fold multiplexed libraries seem to be most common, with between 500,000 and a few million reads dedicated to each individual sample.

In most cases GBS aims at detecting and simultaneously genotyping a large number of single nucleotide polymorphism (SNP) markers. In this study we used GBS on a rice F2 population derived from a cross of an elite cultivar from East Asia (*Oryza sativa* ssp. *Japonica* cv. Nipponbare, NB) and a West African wild rice (*Oryza longistaminata*, OL). Several complex traits are found in OL but are absent in NB. For example, OL is capable of perennial growth, while NB is an annual plant. Furthermore, OL is capable of clonal propagation through the use of rhizomes. To identify the genetic basis of those traits we wanted to perform linkage analysis in an F2 population. Since there are only few markers available for this cross in public datasets and traditional marker development and genotyping can be laborious we established a GBS pipeline.

Performing GBS on an F2 population incurs some specific difficulties since 50 % of all SNP sites are expected to be in a heterozygous state. This demands higher read coverage to accurately call genotypes, since correctly calling a heterozygous allele requires the presence of reads from both allele states (Johnson et al. 2015; Hyma et al. 2015). Some existing GBS pipelines and imputation algorithms deal with that problem by omitting heterozygous calls. In our case that solution was not acceptable, since this would potentially eliminate 50 % of all markers. Another problem associated with using a wild variant in a cross is that there is considerable heterozygosity in the wild parent's genome. This can lead to the inability to correctly infer parental haplotypes. In this F2 population 20 % of all SNP sites found were heterozygous in OL, whereas only 1 % were so in NB. In addition, it might be possible that the wild parent (OL) has genome rearrangements or gene copy number variations as compared to the cultivated parent (NB). Those rearrangements might cause erroneous genotypes in specific regions and linkage of markers which are in reality located on different chromosomes.

By a combination of the comparatively low-cost Illumina MiSeq platform (Loman et al. 2012) and high multiplexing we created a cost-effective, medium throughput (a few hundred to a thousand individuals) genotyping pipeline. The pipeline was designed to specifically address rice F2 populations, but it should be useful for any F2 population. We investigated the effects of two different REs and different levels of multiplexing on the number of detected SNP markers. Also, we provide an example of how relatively low-coverage data (*ca.* 150,000 reads per sample) can be sufficient to generate high density genetic maps. Our pipeline uses simple error correction and imputation methods that take advantage of the long, uniparental haplotype blocks found in F2 populations. To show that our GBS pipeline is producing useful genotypes we mapped QTLs for tiller number and confirmed these QTLs using introgression lines derived from the same parents as the F2 population.

## MATERIALS AND METHODS

### Plant cultivation and population development

The population used in this study was produced and cultivated in the International Rice Research Institute (IRRI), Los Baños, Philippines. An African wild rice, *Oryza longistaminata* Acc. IRGC110404 (OL) as male was crossed with the cultivar *Oryza sativa japonica* cv. Nipponbare (NB) as female to produce F1 plants and subsequently F2 populations by self-pollination. Since NB and OL are rather distant relatives within the *Orzya* genus there is some degree of incompatibility between both parents. Specifically, the cross between NB and OL led to a failure of endosperm development resulting in embryonic death. Therefore embryo-rescue had to be performed to avoid embryonic death ofF1 seeds. In total 301 and 813 F2 plants were grown in the paddy field in the screen house of IRRI in the spring season (Feb-May) and the fall season (Sep-Dec) of 2014, respectively. The total number of tillers (primary and branched shoots of grass plants) was determined after digging up those F2 plants from the paddy field. Leaf blades of the F2s and three replicate individual plants of each, NB and OL were sampled for DNA extraction.

Previously we developed a set of introgression lines (ILs) which harbor between one and three substituted genomic segments derived from OL in the NB genomic background (Ramos et al. 2016). The ILs consist of BC_4_F_7_ and BC_5_F_6_ plants derived from a cross between OL as female and NB as male and successive backcrosses by NB followed by self-fertilization. Four ILs were selected based on the QTL regions found in this study. The ILs and the recurrent parent NB were germinated in a greenhouse and cultivated for 30 days. The seedlings were then transplanted to paddy fields at the research station of Nagoya University, Togo, Aichi Prefecture, Japan. Ten plants per line were planted in each row. The number of tillers was counted at the flowering stage in the ILs and NB excluding damaged plants and plants next to the border of the plot to avoid position effects.

### Library preparation and sequencing

Genomic DNA from plant material was extracted using the cetyltrimethylammonium bromide (CTAB) method (Doyle and Doyle 1987). DNA integrity was analyzed by electrophoresis using a 1 % agarose gel. DNA concentration of each sample was measured using a QuantusTM Fluorometer with a QuantiFluorTM dsDNA system (Promega, Madison, WI, USA) and adjusted to 10 ng μ^−1^. Libraries were prepared using a combination of two restriction enzymes according to (Poland et al. 2012) with the following modifications: Genomic DNA samples (100 ng each) were digested in 20 μl of CutSmart Buffer by 8 units of PstI or *Kpn*I, each with 8 units of *Msp*I (all New England Biolabs (Ipswich, MA, USA), for PstI and *Kpn*I the High-Fidelity version was used). The digestion was performed at 37 °C for 1 h, followed by an inactivation step at 65 °C for 20 min. Ligation was conducted in CutSmart Buffer without any modifications to the original protocol. A set of 192 unique barcodes were selected from the list of 384 barcodes designed for PstI listed in (Poland et al. 2012). These barcodes were utilized for both, adapters with PstI overhang and *Kpn*I overhang. 32-multiplexed libraries for samples digested by PstI and *Msp*I or 96-multiplexed libraries for *Kpn*I and *Msp*I were prepared by pooling samples and subsequent PCR-amplification. DNA qualities and fragment sizes in the prepared libraries were evaluated using a Microchip Electrophoresis System for DNA/RNA analysis (MCE®-202 MultiNA, SHIMADZU, Kyoto, Japan). In total ten 32-multiplexed libraries and nine 96-multiplexed libraries were prepared. The libraries were sequenced using a MiSeq instrument with the MiSeq reagents kit v3 for 150 cycles (Illumina Inc., San Diego, CA, USA).

### Detection of SNPs from raw sequencing data

To detect informative SNPs from raw sequencing data the TASSEL 4 (Trait Analysis by Association, Evolution and Linkage 4) GBS pipeline (Glaubitz et al. 2014) was used. This included creation of a collection of unique, 64 bp long sequences (tags) from the raw sequencing data, alignment of tags to the IRGSP release 7 of the *Oryza sativa* Nipponbare reference genome (Kawahara et al. 2013) using BWA (Burrows-Wheeler Aligner) (Li and Durbin 2009) with the -aln and -samse options, SNP calling and filtering of SNPs based on minor allele frequency. To identify samples with poor read coverage the TASSEL 4 log files for each library were inspected for individuals with very low read coverage (< 1,000 reads in our case). These individuals were removed from the analyses or resequenced in another library if enough plant material was available. We noted that there is a positive correlation between the number of reads and the integrity of the extracted DNA. Initially, SNPs were called without specifying a filter using the DiscoverySNPCallerPlugin from TASSEL 4. Then all SNPs with a minor allele frequency of less than 0.25 were removed, as those likely represented sequencing errors or rare alleles.

In the next step the SNPs were filtered based on parental alleles to leave only SNPs which have fixed, but alternate alleles at any given locus. To achieve this we selected only those SNPs which were: (1) not variable within each set of triplicate parental samples, (2) not heterozygous in either parent and (3) different between both parents. Filtering was performed using the hapmap-formatted files and awk. The resulting collection of SNPs was then thinned out using vcf-tools (Danecek et al. 2011) to a minimum distance of 64 bp between two SNP sites. This eliminated redundant SNPs originating from the same tag, which in most cases had identical parental genotypes within each tag. This collection of SNPs was then used to explore the effects of different levels of missing data and imputation.

Preliminary analyses indicated that the biggest source of error would be undercalled heterozygous alleles (true heterozygous alleles wrongly called as homozygous alleles due to the absence of reads from one of the two states of a heterozygous allele). To counter for this we used vcf-tools to only allow genotypes that are supported by at least 7 reads per site and sample. This limits the probability of undercalling a heterozygous site to a theoretical maximum of 1.6 % (Swarts et al. 2014). In the same step a filter for different levels of missing data was implemented. Specifically, we generated (preimputation, pre-error-correction) datasets in which up to 5 %, 50 % or 75 % of all genotypes for any given site were missing.

### Imputation and error correction

As shown here and in Spindel *et al.* (Spindel et al. 2013) GBS data inherently contains errors and has to be imputed to be useful for linkage analysis. For our work we took advantage of the fact that missing data and wrongly called alleles are randomly distributed across sites and samples. Furthermore, the F2 population in this study is characterized by long-range, uniform parental haplotypes that are long compared to the putative errors. We thus developed a simple imputation and error correction algorithm that is based on regular expressions and executed in R (R Development Core Team 2008).

In the first step the data is transformed from the nucleotide-based hapmap format to an ABH-based format, where A represents NB, B represents OL and H represents heterozygous alleles. After conversion we first imputed missing data. Stretches of missing genotypes were filled with the appropriate allele if both flanking, not missing alleles were of the same state. This imputation resulted in an almost complete elimination of missing alleles. Next, we tried to address the undercalling of heterozygous sites.

Empirically we set a minimum haplotype length of four sites. In any given F2 individual, if a series of homozygous or missing sites of length ≤ 4 was flanked on both sites by a heterozygous allele, this stretch was replaced with heterozygous sites. The other main error type seemed to be single erroneous alleles interspersed in longer homozygous haplotypes. We assumed those errors to come from misalignments of reads, probably due to structural differences in the genome of OL compared to the NB reference genome. To counter for this we used a similar strategy as we used to correct undercalled heterozygous alleles, but used a minimum haplotype length of 1. This procedure reduce the number of missing genotypes as a percentage of all genotypes from 2.07 % to 0.18 % while it increased the number of heterozygous alleles from 46.27 % to 54.57 % (data from the fall 2014 population with up to 75 % missing data per site, full dataset in Table S1). In the final step data from both analyzed populations was combined based on the assumed physical position of SNP markers. Since two different enzymes were used for the spring 2014 and the fall 2014 population no SNP marker was found in both datasets, as different enzymes generate different sets of reads. Thus, we imputed missing data again using the rules devised above to fill in sites. All TASSEL scripts and the scripts used for post-TASSEL data processing can be found in Data S1. The imputation and error correction logic described here (in addition to functions for graphical analyses of genotypes) is also available in the ′ABHgenotypeR’ package for R, which is available at https://github.com/StefanReuscher/ABHgenotvpeR or *via* CRAN (Comprehensive R archive network).

### Data analysis

General data analysis was performed using the TASSEL graphical user interface and R. QTL analyses and simulations were performed using the R package ′qtl’ (v1.37.11) (Broman et al. 2003). For QTL simulations, phenotypic values and genotypes of simulated F2 populations were generated using the function ″sim.cross” implemented in the ‘qtl’ package and described in detail in (Broman and Sen 2009). “sim.cross” requires a genetic map of markers, the number of individuals and a model of QTLs to generate a simulated population. For simulating the genetic map of markers, we used the “sim.map” function which requires chromosome lengths and marker numbers. The lengths of the chromosomes were set to 140 cM, 115 cM, 130 cM, 110 cM, 100 cM, 105 cM, 110 cM, 100 cM, 75 cM, 80 cM, 100 cM and 105 cM for chromosomes 1 to 12, respectively, based on a genetic map of microsatellite markers developed in our previous QTL study for F2 populations derived from a cross between NB and OL. Simulations with 50, 100, 200 or 400 equally spaced markers were performed. For simulating phenotypic values which were affected by a number of simulated QTLs we assumed the existence of eight QTLs (on 8 out of 12 chromosomes), each of which had an additive effect of 0.5. The residual phenotypic variation was assumed to be normally distributed with a variance of 1. Under these assumptions each of the simulated QTLs had 4.17 % contribution to the phenotypic variance. With the simulated genetic map and the QTL model, data sets of F2 populations for 200, 400, 600, 800 and 1000 individuals were generated using ″sim.cross”. We performed simple interval mapping in the simulated F2 populations using the function ″scanone” with the multiple imputation method (Sen and Churchill 2001). In the multiple imputation method genotypes between markers were imputed with 1 cM intervals based on genotypes of flanking markers and multiple imputed genotype data were generated for each individual. Then, a linear regression model was fitted for each marker using the imputed genotype data and the phenotype data with the assumption of normal distribution of phenotypic values. The threshold for significant LOD scores was calculated from 1,000 permutation tests. According to past studies, confidence intervals of detected QTLs were usually larger than 10 cM (Darvasi 1998; Kearsey and Farquhar 1998), so we used that size as a threshold. If a significant QTL (P ≤ 0.05) was detected around the simulated, true QTL position (±10 cM), we counted it as correctly detected. For each condition 100 simulations were performed and the probability to correctly detect all QTLs was calculated.

Genetic maps using real data were constructed using the ″est.map” function with default parameters. QTL analyses for the number of tillers in 1,081 F2 plants was performed using a linear regression model with the multiple imputation method by ″scanone”. The threshold for significant LOD scores was calculated from 1,000 permutation tests. The 95 % confidence intervals of significant QTLs were estimated using the function “bayesint” which takes 10^LODscore^ values for an obtained LOD profile and rescales it to have an area of 1, followed by calculating the connected interval having 95 % coverage of the area. The function ″fitqtl” was used for calculating percentages of variance of the significant QTLs by calculating the coefficient of determination for each single-QTL model obtained using ″scanone”. Additive and dominant effects of the significant QTLs were calculated from mean phenotypic values for each genotype at the QTL positions obtained by using the function ″effectplot”.

Genome-wide analysis of restriction sites were performed using the ″restric” tool from the emboss software suite (Rice et al. 2000). Random sampling of reads from fastq files was performed using fastq-tools (http://homes.cs.washington.edu/~dciones/fastq-tools/).

### Statement on data availability

Dataset SI contains all code necessary to replicate the GBS-pipeline. The data imputation and error-correction logic is also available as the R package “ABHgenotypeR”. Dataset S2 contains all genotypes from this study, including marker order and position. Complete genotype and SNP descriptions are available upon request.

## RESULTS

### Application of GBS to a rice F2 population

A population of 268 F2 plants from a cross of NB and OL, including triplicate parental samples, from the spring 2014 season was sequenced first. From this population libraries of 32 samples each were prepared and processed with the GBS pipeline (Fig. 1). This approach resulted in 618,844 average reads per individual, which yielded 108,905 potentially useful SNP sites before the application of any filtering (Table 1). Filtering those sites resulted in at least 2,144 markers (5 % missing data). Analyses using simulated data to determine QTL detection probabilities showed that this number of markers is more than sufficient to detect even weak QTL (see Figure S1). In fact, a few hundred markers gave sufficient detection power while at the same time the number of F2 individuals appears to be the limiting factor. We therefore optimized our GBS pipeline to process more F2 individuals at the expense of generating a lower number of SNP markers by multiplexing more samples per library.

**Figure 1:**
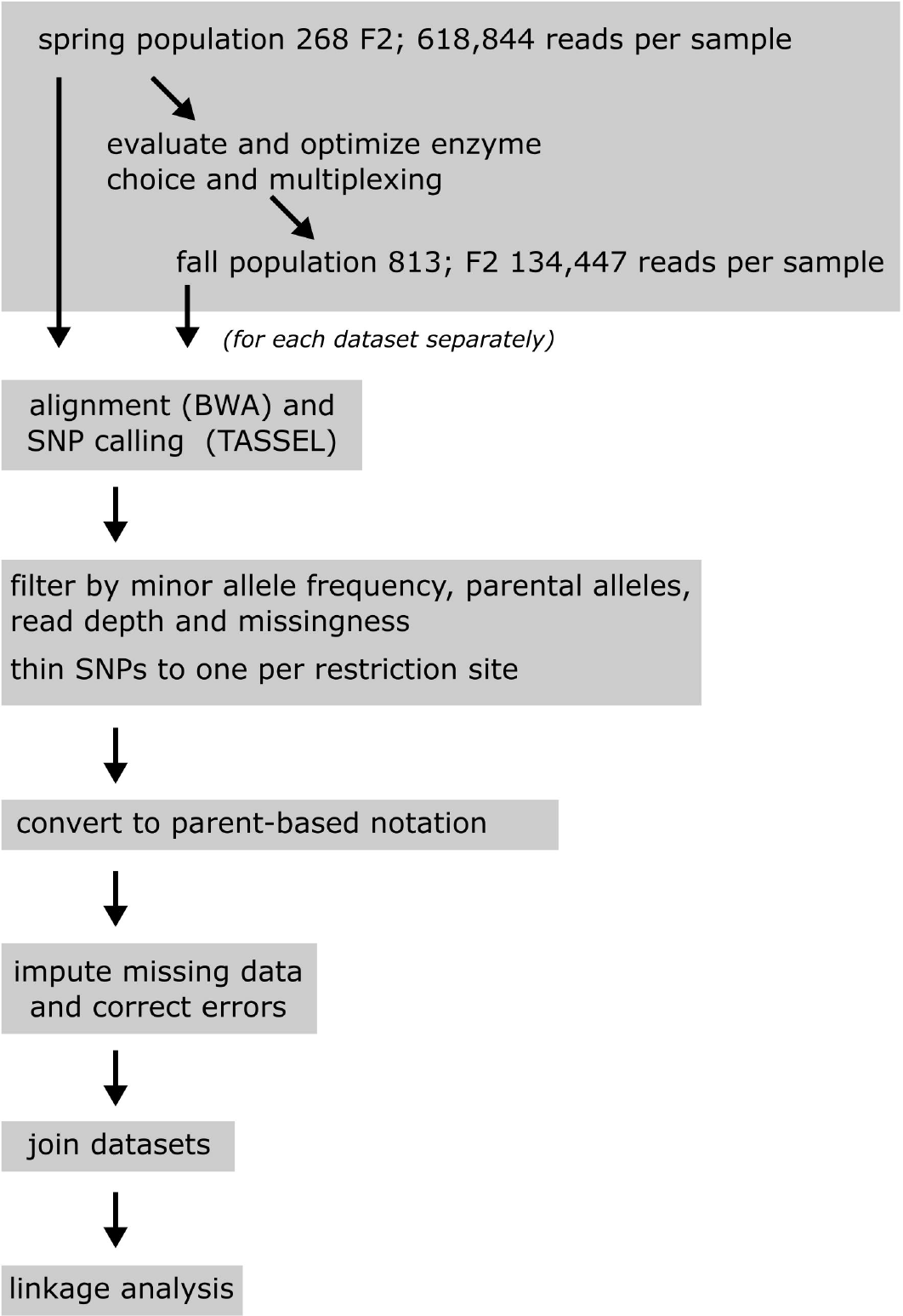
**Flowchart of the GBS data processing** A schematic overview of the different steps of the GBS pipeline.

**Table 1.** **Basic parameters of both GBS experiments described in this work.**

|  | spring 2014 | fall 2014 |
| --- | --- | --- |
| enzymes | *PstI-MspI* | *KpnI-MspI* |
| number of F2 individuals | 268 | 813 |
| multiplexing | 32 | 96 |
| reads per sample (mean ± SD)<sup>a</sup> | 618,843.7 ± 178,135.8 | 134,447.3 ± 50,788.87 |
| no. of initial SNPs<sup>b</sup> | 108,905 | 37,938 |
| no. of sites (< 5% missing data) | 2,144 | 301 |
| no. of sites (< 50% missing data) | 5,812 | 837 |
| no. of sites (< 75% missing data) | 7,058 | 1,096 |

<sup>a</sup> Numbers are based on good, barcoded, aligned reads.
<sup>b</sup> Number of SNP sites called by TASSEL before any filtering steps were applied.

For a larger population of 813 F2 plants and triplicate parental samples from the fall 2014 season the following changes were implemented: (1) Instead of using PstI as the rare-cutting enzyme we used *Kpn*I. There are 107,953 PstI cut sites reported in the NB reference genome, while there are only 45,065 *Kpn*I cut sites. Thus, if all parameters were kept constant, in libraries prepared with *Kpn*I the resulting reads will be distributed among fewer sites, but reach a higher per-site coverage. (2) Taking advantage of the higher per-site coverage using *Kpn*I we increased the number of samples per library. Prior to library preparation we examined the effects of decreased read coverage per F2 individual by randomly sampling a fraction of reads from each input fastq file. In these simulated multiplexing analyses it became clear that the undercalling of heterozygous sites (50 % in an F2 population) would become a large source of errors if multiplexing is increased (see Figure S2). Based on those results 96-fold multiplexing was deemed feasible and implemented with the fall 2014 population. This resulted in an average of 134,447 reads per F2 which yielded 37,938 potential SNP sites. After processing and allowing up to 5 % missing data a minimum of 301 SNP sites remained for analysis.

As expected, higher multiplexing and a change to *Kpn*I led to a lower number of detected SNP sites. When processed through our GBS pipeline however, both datasets led to similar genotype patterns, the main difference being the number of sites that were reliably detected. As the final step of the GBS pipeline both datasets were merged. To describe and evaluate the results of the GBS pipeline we subsequently use data from the fall 2014 dataset. For results regarding the genetics of the NB x OL F2 population and linkage analysis we used the combined datasets to maximize detection power and resolution.

### Analysis of general SNP properties

The unfiltered GBS dataset contained a high proportion of missing data (Fig. 2A) and only ca. 4,500 out of 37,938 sites were detected in all samples. Also, a substantial number of SNPs was observed with very low minor allele frequencies (MAF) (Fig. 2B). We used a threshold of MAF > 0.25 and different proportions of missing data (< 5 %, < 50 %, < 75 %) and analyzed the MAF and the proportion of heterozygous sites. When using a very stringent filter of < 5 % missing data both, the MAF and the proportion of heterozygous sites reached a lower limit at around 0.35 (Fig. 2C and D). At a higher proportion of missing data some sites could be observed which had a MAF and proportion of heterozygosity as low as the set threshold of 0.25 (Fig. 2 E to H). The bigger spread in allele frequencies and heterozygosity observed for datasets with a higher percentage of missing data might be explained by the inclusion of sites with low read coverage in those datasets. SNP sites which are supported by a small number of reads are more prone to errors. For example, reads representing either NB or OL alleles could have different amplification efficiencies during library preparation. For SNPs with high read coverage this might have no effects, but for SNPs with low read coverage this might skew our ability to detect a specific allele. This observation highlights the importance of both, adequate read coverage and post SNP-calling error correction.

**Figure 2:**
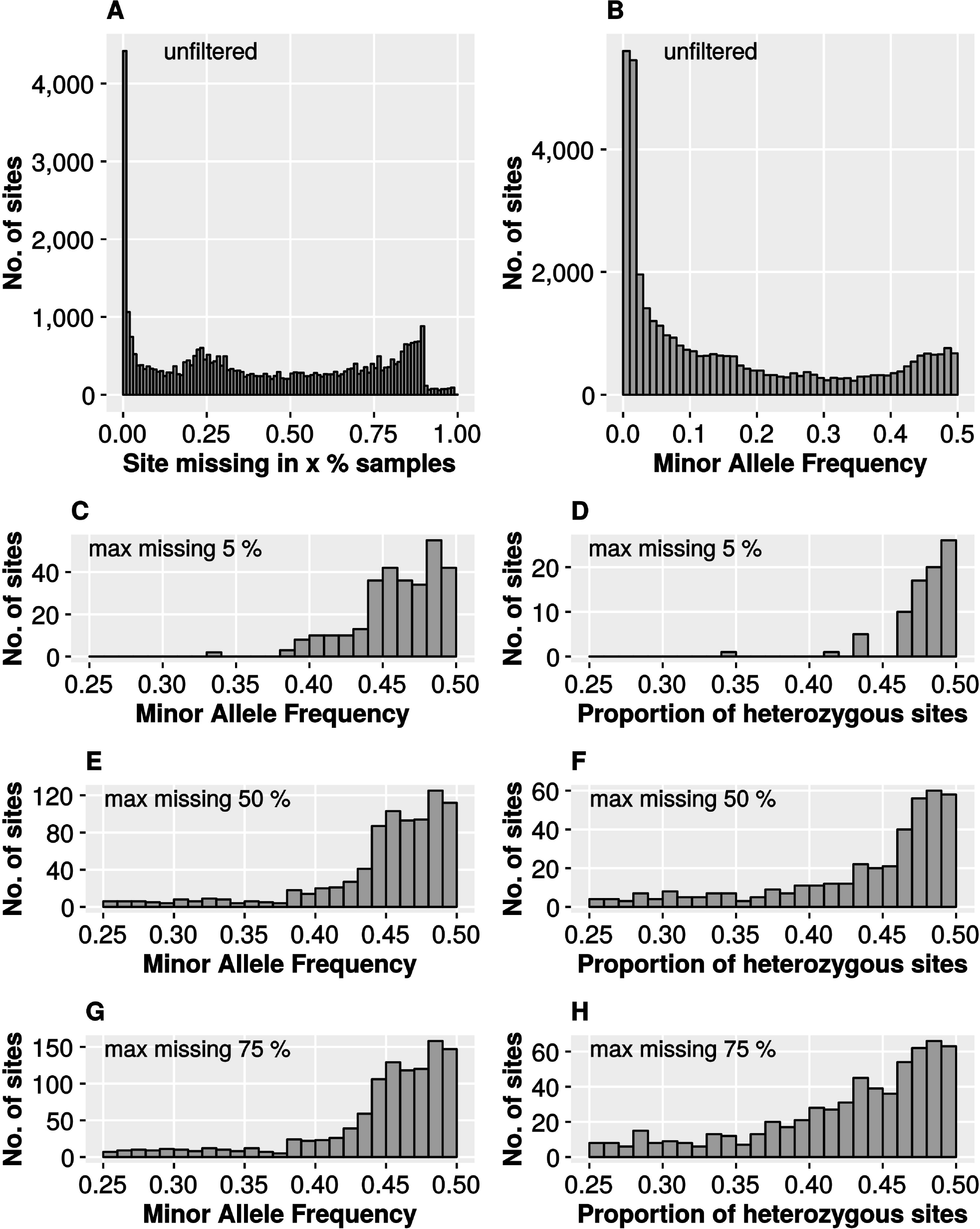
**Basic SNP characteristics using different filter settings for missing data.** Shown are histograms representing the number of SNP sites that exhibit a certain sample coverage (A), minor allele frequency (B, C, E, G) or proportion of heterozygous sites (D, F, H). Data is from 813 F2 plants from the fall 2014 population and was generated using the TASSEL 4 site report function. For A and B unfiltered data directly after SNP calling was used. For C to H, SNP sites were filtered by the indicated proportion of missing data per sample, but no further data imputation or error correction was performed.

To evaluate the fidelity of GBS genotypes we independently genotyped 93 F2 plants using simple-sequence repeat (SSR) markers and compared both sets of genotypes. It was found that the majority of parental genotypes > 90 %) was identical when the results of both genotyping systems were compared (see Figure S3). The 10 % disagreeing markers are explained by single SSR markers in which up to 1/3 of all genotypes disagree and probably by the SSR marker and the closest GBS marker being on different sides of a recombination event.

Next we evaluated the distribution of SNP sites along the chromosome (Fig. 3). SNP sites were notably sparser in the centromeric regions, probably as a result of a high amount of repetitive sequence elements which prevent reads to be mapped to a unique position. Also the distribution of sites along the chromosome arms was not even. In general, the SNP density at any given chromosome position increased with the amount of missing data allowed. However, there were some chromosomal region with low SNP density in which the number of SNPs was hardly affected by the amount of missing data. This was not caused by uneven distribution of *Kpn*I recognition sites (data no shown). For example, a SNP density below the average was observed on the long arms of chromosome 4 and chromosome 9. The occurrence of such SNP deserts was observed before (Wang et al. 2009; Krishnan S et al. 2014), but it is unclear if and how those regions are associated with domestication.

**Figure 3:**
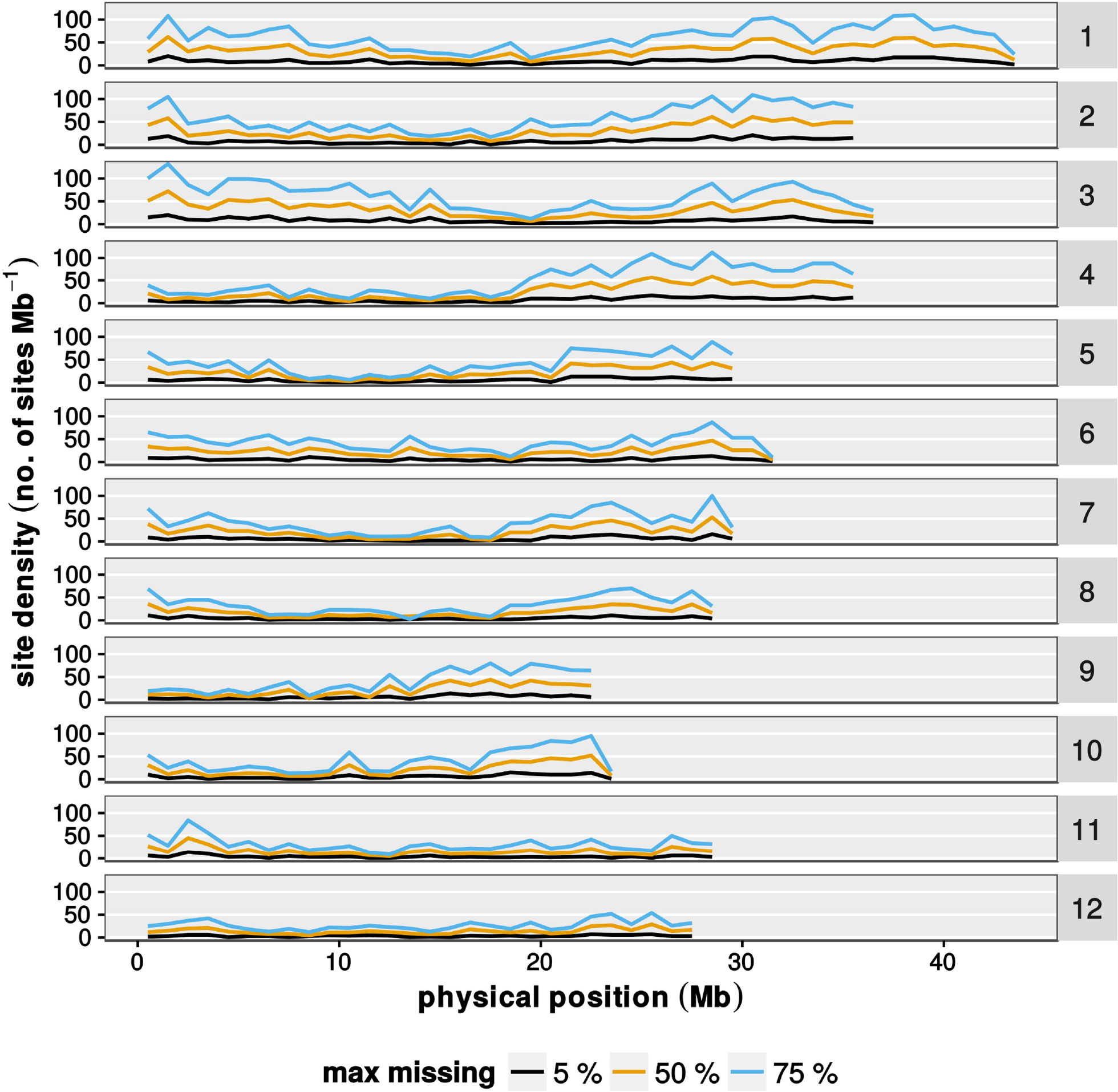
**Marker densities along the chromosomes.** Shown is the marker density along the 12 rice chromosomes. The number of markers was determined for bins of 1 Mb. Different colored lines represent datasets with the indicated proportion of missing data. Data is from the fall 2014 population (n = 813 individuals).

In an ideal F2 population one would expect that the parental alleles segregate according to a 1:2:1 ratio (parent A : heterozygous : parent B). However, a plot of allele states along the chromosomes revealed regions with distorted genotype ratios (Fig. 4). As a general trend the OL alleles seemed to be transmitted at slightly lower levels. As an extreme example, the long arm of chromosome 4 has a drastically reduced frequency of the OL allele, with OL genotype frequencies decreasing to less than 10 %, as opposed to the expected 25 %. In most chromosomal regions where one parental allele was found underrepresented the frequency of heterozygous genotypes in turn was increased to more than 50 %. Very likely those effects are due to chromosomal regions associated with reproductive incompatibility.

**Figure 4:**
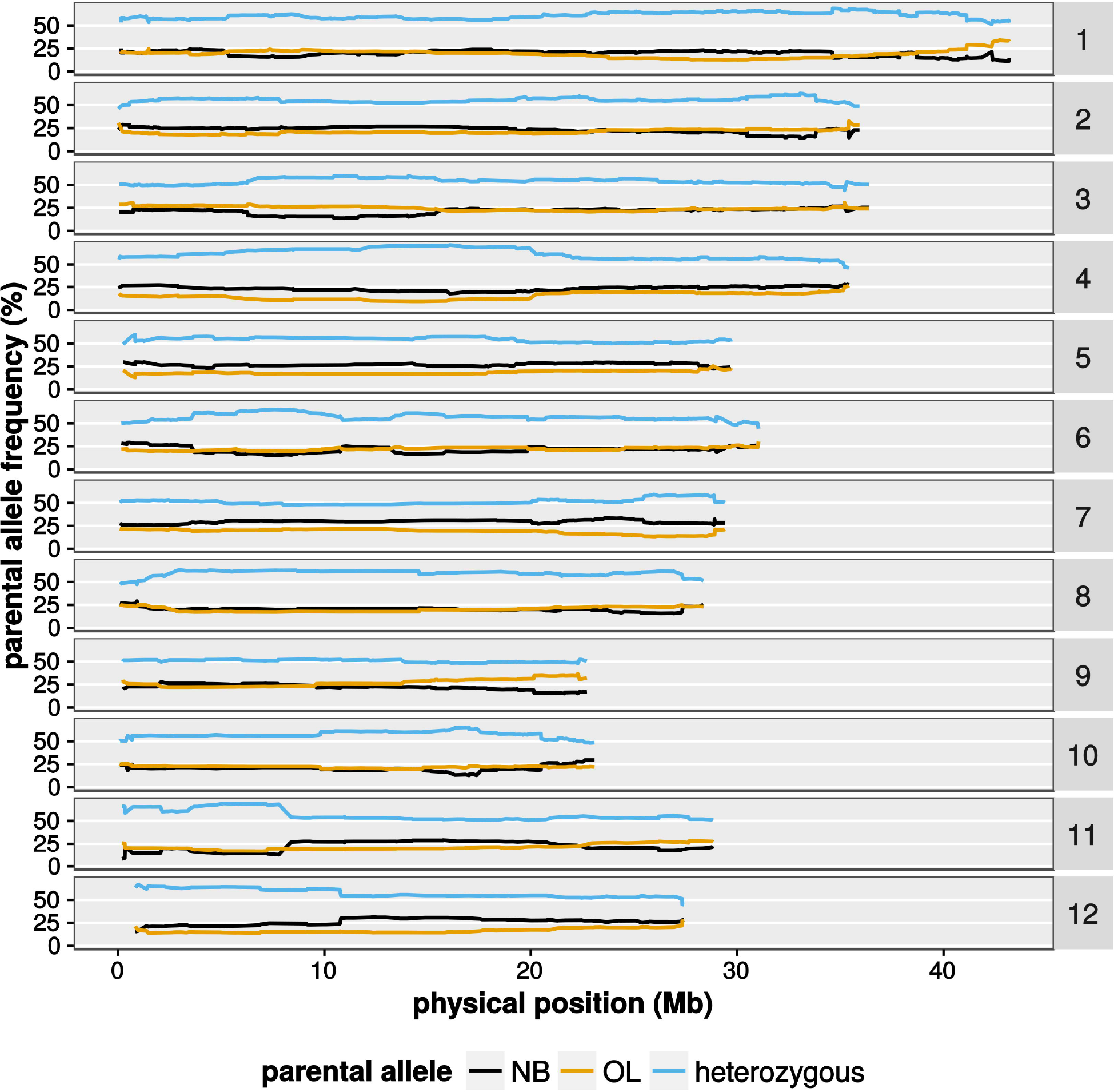
**Parental allele frequencies along the chromosomes.** Shown are the frequencies of parental alleles observed along the 12 rice chromosomes. Data is from the joined datasets from spring and fall 2014. Only marker present in 95 % of all samples in the respective dataset are shown.

### Constructing a genetic map

To inspect GBS genotypes and haplotypes we constructed graphical representations of genotypes (Fig. 5, full dataset in Data S2). This made it obvious that GBS data without imputation and error correction contains wrongly called genotypes (Fig. 5A). Since F2 populations have relatively long haplotypes the observed very short (1-2 markers) uniform genotype stretches found as islands in longer stretches are most likely errors. After imputation of missing data (Fig. 5B) we used a simple error correction algorithm based on haplotype length to efficiently correct those errors (Fig. 5C).

**Figure 5:**
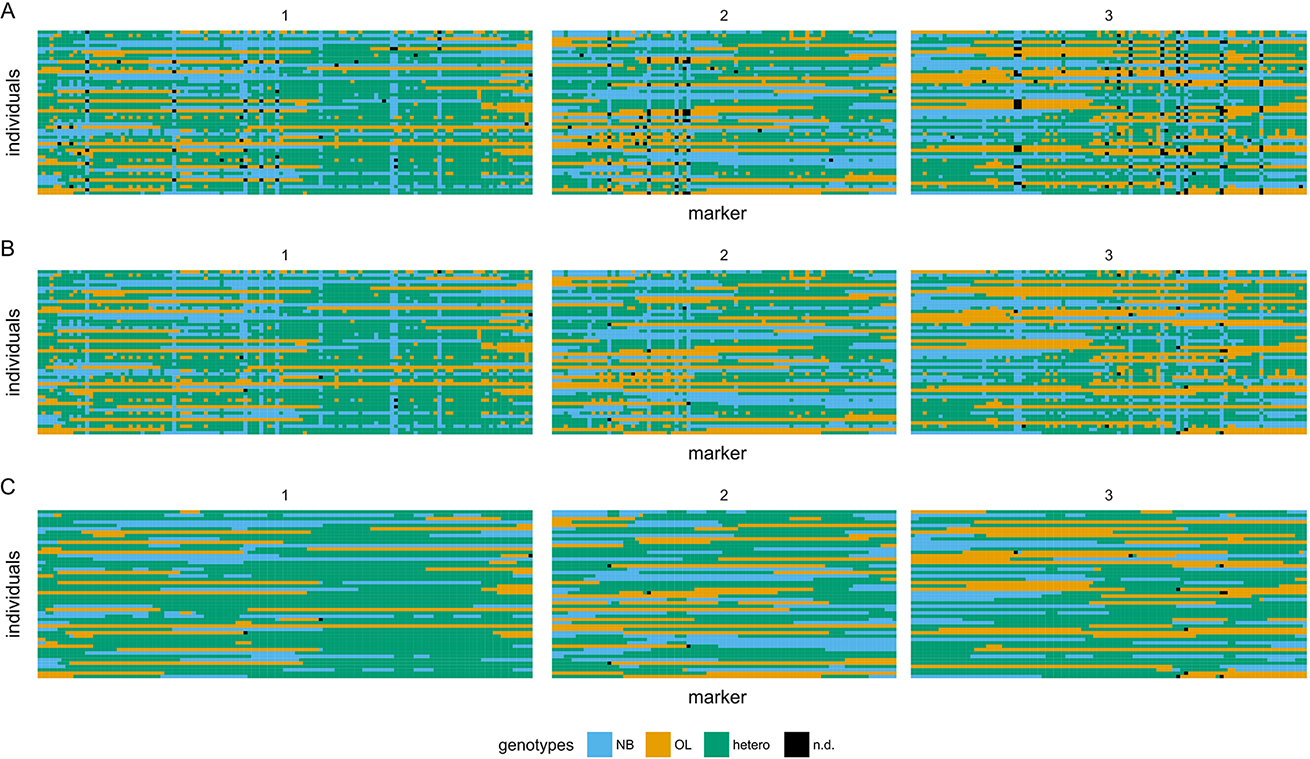
**Graphical representations of GBS-derived genotypes at different stages of post-processing.** Shown are graphical representations of genotypes after inferring parental alleles (A), after inferring parental alleles and imputation of missing data (B) and after inferring of parental alleles, imputation and error correction (C). Genotypes of 50 representative F2 individuals are shown, with each F2 as a single horizontal track. The chromosome length is proportional to the number of markers and only chromosome 1 to 3 are shown. In total 312 markers (fall 2014 population, up to 50 % missing data) are displayed with genotypes color-coded as blue (NB), orange (OL), green (heterozygous) and black (not determined).

When we used the fall 2014 dataset to construct a genetic map it became again clear that raw GBS data cannot be used directly (Fig. 6). When uncorrected data with up to 75 % or 50 % (Fig. 6 A; B, D and E) of missing data per site was used to generate a genetic map, chromosomes appeared expanded with chromosomes of up to 3,500 cM. The map distention we observed was conspicuously similar to the distention shown in (Spindel et al. 2013) and we applied a similar strategy to consolidate our genetic map. Both, a rigorous restriction on missing data (up to 5 % missing, Fig. 6 G–I) or imputation and error correction (Fig. 6 C and F) seemed to alleviate the problem. Restricting missing data led to a strong reduction of available SNP sites (compare 837 for 50 % missing to 301 for 5 % missing) but also shortened the genetic map. Using filtering, imputation and error correction we gradually improved the genetic map even when up to 75% of genotypes were initially missing for each individual site. The final genetic map (Fig. 6 I) had a total size of 1,536 cM which is in agreement with other data. We still observed some distention, for example on chromosome 5 and chromosome 12. Although haplotypes and alleles appear to be correct in those region we can observe strong linkage of markers in those region with markers from different chromosomes (data not shown).

**Figure 6:**
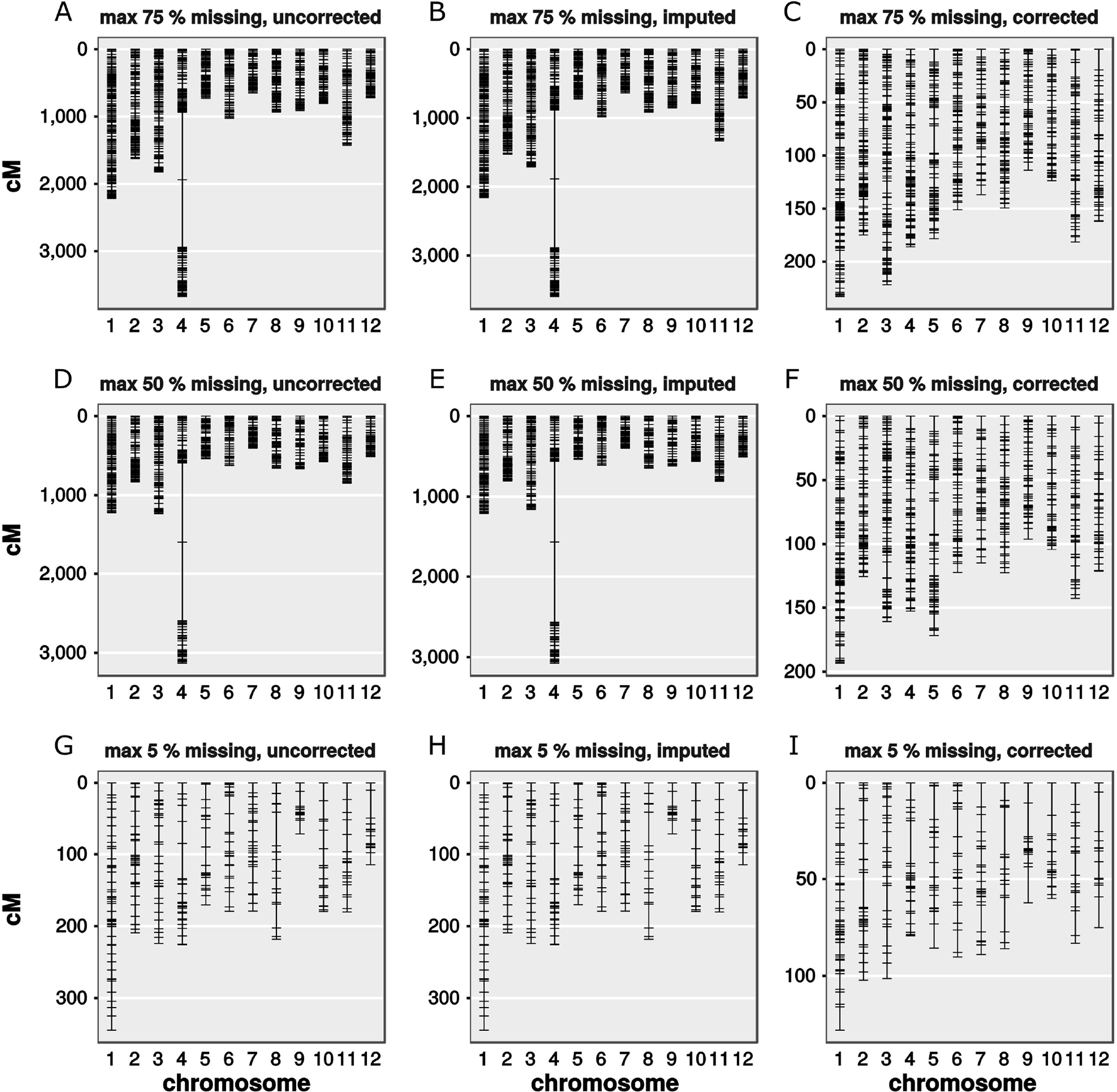
**Genetic maps from datasets with different proportions of missing data and post-processing.** Shown are linkage maps of GBS marker datasets. Panels show datasets with SNP-calling thresholds allowing up to 75 % (A-C), 50 % (D-F) and 5 % (G-I) missing data, at different steps of the GBS pipeline. Uncorrected (A, D, G) indicates data without further post-processing. Imputed (B, E, H) indicates data with missing data imputed, but no error correction performed. Corrected (C, F, I) indicates data with both, imputation and error-correction performed. Data is from 813 F2 plants from the fall 2014 dataset. Distances between markers are shown in centimorgan (cM).

### QTL analysis

Being able to produce a correct genetic map using the combined dataset reassured us that our GBS data is sufficient for linkage analysis. For QTL analysis in 1,081 F2 plants we chose to use the number of tiller as the phenotype. We detected four significant QTLs on chromosomes 1, 3, 4 and 8 which were named qOLTN1, qOLTN2, qOLTN3 and qOLTN4, respectively (Fig. 7A). Among these four QTLs qOLTN1 on chromosome 1 showed the highest LOD score with 20.15 (Fig. 7B), while the other QTLs showed LOD scores less than 6.9. To analyze these QTLs in more detail, we calculated 95 % confidence intervals, percentages of variance and effects for each QTL (Table 2). The confidence interval of qOLTN1 spanned a 3.6 Mb region from 27. 1 Mb to 30.7 Mb on chromosome 1. This QTL explained 8.23 % of the variance in the number of tillers of the F2 population and showed a negative additive effect of −9.17 and a positive dominant effect of 5.22. These results suggested that an OL allele at qOLTN1 acts recessive to decrease the number of tillers as compared to NB. Unlike the case of qOLTN1, the other QTLs gave only little contributions on the differences in the number of tillers and relatively smaller effects (Table 2). Interestingly, qOLTN4 exhibited a positive superdominant effect in which the additive effect was −2.44 while the dominant effect was 4.51. This result means that heterozygotes at qOLTN4 produce more tillers than either NB homozygotes or OL homozygotes.

**Figure 7:**
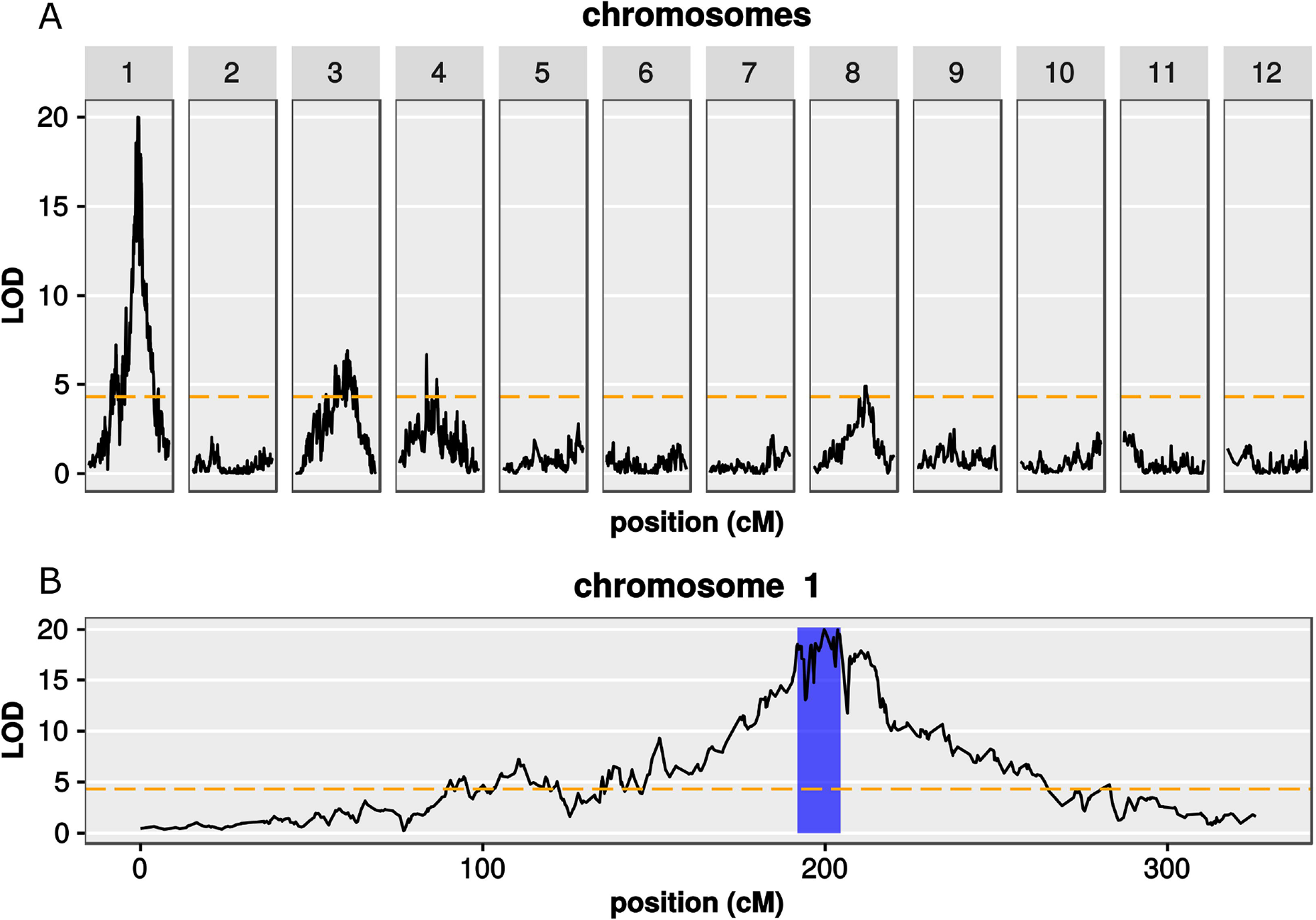
**Detection of QTL for tiller number using GBS markers.** Shown are the results of a linkage analysis to detect QTL that have an effect on tiller number using data from joined spring and fall datasets with up to 75 % missing data per marker. LOD scores are shown as black lines for all 12 chromosomes (A) or for chromosome 1 only (B). A LOD threshold for significance (P ≤ 0.05) is shown as a dashed orange line. The blue area in (B) highlights the 95 % confidence interval of qOLTN1 (QTL1 for tiller number *Oryza longistaminata)*. Distances are shown in centimorgan (cM).

**Table 2.**
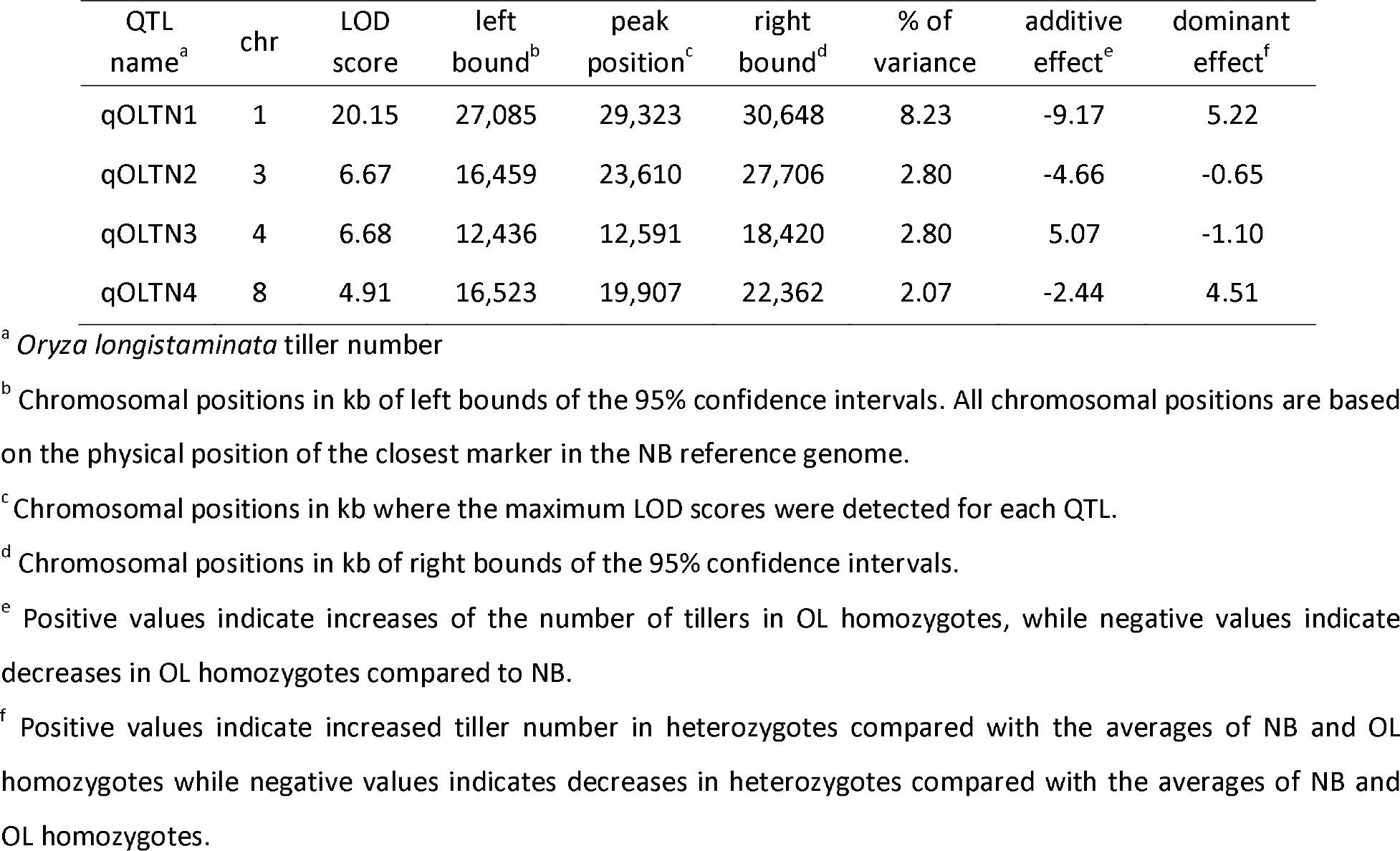
**Percentages of variance and effects of the significant QTLs.**

| QTL name<sup>a</sup> | chr | LOD score | left bound<sup>b</sup> | peak position<sup>c</sup> | right bound<sup>d</sup> | % of variance | additive effect<sup>e</sup> | dominant effect<sup>f</sup> |
| --- | --- | --- | --- | --- | --- | --- | --- | --- |
| qOLTN1 | 1 | 20.15 | 27,085 | 29,323 | 30,648 | 8.23 | -9.17 | 5.22 |
| qOLTN2 | 3 | 6.67 | 16,459 | 23,610 | 27,706 | 2.80 | -4.66 | -0.65 |
| qOLTN3 | 4 | 6.68 | 12,436 | 12,591 | 18,420 | 2.80 | 5.07 | -1.10 |
| qOLTN4 | 8 | 4.91 | 16,523 | 19,907 | 22,362 | 2.07 | -2.44 | 4.51 |

<sup>a</sup> *Oryza longistaminata* tiller number
<sup>b</sup> Chromosomal positions in kb of left bounds of the 95% confidence intervals. All chromosomal positions are based on the physical position of the closest marker in the NB reference genome.
<sup>c</sup> Chromosomal positions in kb where the maximum LOD scores were detected for each QTL.
<sup>d</sup> Chromosomal positions in kb of right bounds of the 95% confidence intervals.
<sup>e</sup> Positive values indicate increases of the number of tillers in OL homozygotes, while negative values indicate decreases in OL homozygotes compared to NB.
<sup>f</sup> Positive values indicate increased tiller number in heterozygotes compared with the averages of NB and OL homozygotes while negative values indicates decreases in heterozygotes compared with the averages of NB and OL homozygotes.

To evaluate the results of our QTL simulation (see Figure S1) against this real data we performed linkage analyses for random subsets of varying numbers of F2 plants. As predicted in our simulations, we found that up to 1,000 F2 plants are necessary to reliably detect all significant QTL (see Figure S4). When we used all 1081 F2 plants for linkage analysis but varied the amount of missing data allowed in the prefiltered datasets we found very similar LOD score profiles (see Figure S5). We thus used the dataset with up to 75 % missing data per site before post-processing to maximize marker resolution.

To verify the QTLs we conducted a field experiment to measure the number of tillers in introgression lines (ILs) having OL genomic segments at each of the QTL locations. Four ILs having OL chromosomal segments around QTL locations were selected from the pool of ILs and named IL-qOLTN1, IL-qOLTN2, IL-qOLTN3 and IL-qOLTN4 for having OL chromosomal segments around qOLTN1, qOLTN2, qOLTN3 and qOLTN4, respectively. IL-qOLTN1 and IL-qOLTN2 showed a significant decrease in the number of tillers compared with NB (Table 3). The reductions of tillers in these two ILs is in agreement with the negative additive effects of qOLTN1 and qOLTN2 (Table 2). Furthermore, IL-qOLTN3 and IL-qOLTN4 produced more and less tillers than NB, although the differences were not significant. However, the results observed in IL-qOLTN3 and IL-qOLTN4 also corresponded to the positive and negative additive effects of qOLTN3 and qOLTN4, respectively. In summary, we could successfully detect four QTLs using our GBS data for the number of tillers and verify the effects of those QTLs in ILs.

**Table 3.** **Tiller number in the introgression lines.**

| genotype<sup>a</sup> | chr.<sup>b</sup> | markers<sup>c</sup> | position<sup>d</sup> | No. of tillers<sup>e</sup> |
| --- | --- | --- | --- | --- |
| NB | - | - | - | 13.67 ± 1.70 |
| IL-qOLTN1 | 1 | RM1287-RM297 | 10.8-33.8 | 8.14 ± 1.64 * |
| IL-qOLTN2 | 3 | OL3L26-RM3436 | 5.4-28.2 | 11.13 ± 1.27 * |
| IL-qOLTN3 | 4 | End-RM3866 | 0-23.8 | 15.00 ± 3.12 |
| IL-qOLTN4 | 8 | RM1235-RM5485 | 12.1-24.2 | 11.67 ± 2.87 |

<sup>a</sup> NB indicates Nipponbare, IL-qOLTN1 to 4 indicates introgression lines that carry the respective QTL for tiller number
<sup>b</sup> Chromosome which have an *O. longistaminata* chromosomal segment.
<sup>c</sup> Flanking simple sequence repeat markers of an introgressed *O. longistaminata* chromosomal segment. "End" indicates the end of short arm.
<sup>d</sup> Physical positions of the flanking SSR markers in Mb.
<sup>e</sup> The number of tillers measured in 7-9 plants for each line are shown in mean ± sd. \* indicates a significant difference compared with NB at $P \leq 0.05$ according to Student's t-test with Bonferroni-Holm correction for multiple testing.

## DISCUSSION

Our aim for this study was to utilize GBS for rapid genotyping of rice F2 populations. As expected, GBS proved to be a robust and efficient method to genotype large populations (Elshire et al. 2011; Lu et al. 2013; Spindel et al. 2013; Liu et al. 2014). For successful application of GBS it is necessary to generate adequate read coverage across the genome and also for each individual that is sequenced. In our approach to genotype a rice F2 population we further took into account the number of individuals and markers that are necessary to detect QTLs. Since one of the main motivations to perform GBS is to save time and money compared to classical markers one would like to use as few sequencing runs on any platform as necessary to achieve the desired sequencing depth. Our choice to change the enzyme from *Pst*I to *Kpn*I lead to predictable changes in the resulting SNP collection. Also other reports showed that enzyme choice is an important parameter to optimize GBS for any given species (Heffelfinger et al. 2014). Marker density depends also partially on sequencing depth, which in turn depends on the number of individual per sequencing run. To be most efficient it is thus advisable to take into account the desired marker density when laying out a genotyping project involving GBS. In our experience, performing a small-scale pilot experiment using the desired population and sequencing platform, combined with linkage analysis on simulated data, allowed us to use GBS more efficiently. The results of linkage analyses using both, simulated (see Figure S1) and experimental data (see Figures S4 and S5) suggested that our GBS approach resulted in a saturation of markers. The fact that our linkage analysis yields comparable results even when up to 75 % of missing data for each marker were acceptable in the raw data shows that even simple imputation algorithms can reinforce the usefulness of GBS data tremendously. We speculate that for certain applications even less markers and in turn less reads per individual would be sufficient, thus allowing even higher multiplexing and sample throughput. Of course, this might also depend on the genome size of the analyzed species and the amount of repetitive elements in that genome.

### Optimizing GBS strategies

Although we were successful in using GBS we noticed several shortcomings where no best practice seems to be established yet. There seems to be very little consensus about how GBS protocols should be adapted to different species and to different populations. For variant calling, filtering and exploration of our dataset we used the TASSEL4 (Glaubitz et al. 2014) which was develop to work efficiently with large maize populations. It became apparent that additional specific bioinformatics analyses were necessary to get the most information from our dataset. This shows that a given GBS protocol needs to be optimized for a specific species or population. Another issue is the high error rate of raw GBS data. While it is possible to eliminate most errors using post SNP-calling error correction some errors will inevitably remain. It would be worth to investigate the source of some errors, as this might lead to new insights into the population in question. In our case, where a wild species is crossed to a cultivar it can be assumed that there will be large-scale differences between the two parental genome contributing to the F2 individuals. Those differences most likely include gene copy number variations or even rearrangements of regions between chromosomes. Since we only have a reference genome available for one of the parents at the moment we have no way to control directly for those potential sources of errors. We found indirect evidence for such large rearrangements when we looked at genome-wide linkage of markers. We found several regions in which seemingly correct haplotypes were in strong linkage disequilibrium with both, neighboring regions and regions on other chromosomes (data not shown). Future GBS pipelines could address those issues, either by taking into account improved reference genome information or through linkage disequilibrium filtering.

When we established our GBS pipeline we noticed several irregularities in the genome-wide SNP statistics. For example, we noticed that several regions of the genome were sparsely covered with SNPs (Fig. 3). Also we noted that in several region the parental allele frequency was deviating from the expected 1:2:1 ratio (Fig. 4). It is important to note that this population is affected by reproductive incompatibilities and we had to routinely use embryo-rescue to propagate plant materials. It is very likely that the deviating allele frequency is a consequence of reproductive incompatibility which has its genetic basis in these regions. To further analyze this it would be necessary to genotype the offspring of multiple F1 crosses. We suggest that GBS might be a useful tool to study reproductive isolation and preferential transmission, since it can quickly define regions with allele distortion.

## CONCLUSION

In summary we show an application of GBS to perform linkage analysis in a rice F2 population. We also provide an example on how to plan and carry out adequate, cost effective reduced-representation sequencing. With our dataset we successfully detected QTLs for tiller number on chromosomes 1, 3, 4 and 8 which we could confirm using ILs. We suggest for future GBS genotyping efforts to evaluate enzyme choice, multiplexing of libraries and post-processing to meet the requirements of the desired post-GBS analyses. We predict that the efficiency of GBS in terms of pricing and time will improve even more in the future.

## LIST OF ABBREVIATIONS

GBS: Genotyping-by-sequencing
NB: *Oryza sativa japonica* ′Nipponbare’
OL: *Oryza longistaminata*
RE: restriction enzyme
MAF: minor allele frequency
IL: introgression lines
SNMP: single-nucleotide polymorphism
SSR: simple sequence repeat
BWA: Burrows-Wheeler Aligner
QTL: quantitative trait locus
GWAS: genome-wide association study
TASSEL: trait analysis by association evolution and linkage.

## AUTHORS’ CONTRIBUTIONS

TF created the NB × OL F2 populations, prepared sequencing libraries, performed MiSeq sequencing, performed linkage analysis and assisted in drafting the manuscript. SR developed and optimized the GBS pipeline, performed bioinformatics analyses and drafted the manuscript. KD and MA conceived the experimental design. KKJ oversaw the production of plant materials. MA oversaw the project and assisted in drafting the manuscript. All authors have read and approved the manuscript.

## ACKNOWLEDGMENTS

The authors like to thank Dr. Rosalyn B. Angeles-Shim, Ruby S. Lapis and Angelito A. Aragon from the International Rice Research Institute for cultivating our plant material and Yoko Niimi for performing genotyping using SSR marker. This work was supported by Core Research for Evolutional Science and Technology from the Japan Science and Technology Agency to MA, the Council for Science, Technology and Innovation, Cross-ministerial Strategic Innovation Promotion Program, Technologies for creating next-generation agriculture, forestry and fisheries (funding agency Bio-oriented Technology Research Advancement Institution, NARO) to KD and the National Bio Resource Project (NBRP) “Rice” to KD

## REFERENCES

Begum,H., Spindel,J. E., Lalusin,A., Borromeo,T., Gregorio,G. et al., 2015 Genome-wide association mapping for yield and other agronomic traits in an elite breeding population of tropical rice (Oryza sativa). PLoS ONE 10: e0119873.

Broman,K. W., Wu,H., Sen,S., and Churchill,G. A., 2003 R/qtl: QTL mapping in experimental crosses. Bioinformatics 19: 889–890.

Broman,K. W., and Sen,Ś., 2009 A guide to QTL mapping with R/qtl Springer, New York.

Burrell,A. M., Pepper,A. E., Hodnett,G., Goolsby,J. A., Overholt,W. A. et al., 2015 Exploring origins, invasion history and genetic diversity of Imperata cylindrica (L.) P. Beauv. (Cogongrass) in the United States using genotyping by sequencing. Mol Ecol 24: 2177–2193.

Danecek,P., Auton,A., Abecasis,G., Albers,C. A., Banks,E. et al., 2011 The variant call format and VCFtools. Bioinformatics 27: 2156–2158.

Darvasi,A., 1998 Experimental strategies for the genetic dissection of complex traits in animal models. Nature Genet 18:19–24.

Davey,J. W., Hohenlohe,P. A., Etter,P. D., Boone,J. Q., Catchen,J. M. *etai*, 2011 Genome-wide genetic marker discovery and genotyping using next-generation sequencing. Nat Rev Genet 12: 499–510.

Donato,M. de, Peters,S. O., Mitchell,S. E., Hussain,T., and Imumorin,I. G., 2013 Genotyping-by-sequencing (GBS): a novel, efficient and cost-effective genotyping method for cattle using next-generation sequencing. PLoS ONE 8: e62137.

Doyle,J. J., and Doyle,J. L., 1987 A rapid DNA isolation procedure for small quantities of fresh leaf tissue. Phytochemical Bulletin: 11–15.

Duitama,J., Silva,A., Sanabria,Y., Cruz,D. F., Quintero,C. et al., 2015 Whole genome sequencing of elite rice cultivars as a comprehensive information resource for marker assisted selection. PLoS ONE 10: e0124617.

Elmer,I., Humira,S., and FranÇois,B., 2015 Association mapping of QTLs for sclerotinia stem Rot resistance in a collection of soybean plant introductions using a genotyping by sequencing (GBS) approach. BMC Plant Biol 15: 5.

Elshire,R. J., Glaubitz,J. C., Sun,Q., Poland,J. A., Kawamoto,K. et al., 2011 A robust, simple genotyping-by-sequencing (GBS) approach for high diversity species. PLoS ONE 6: el9379.

Glaubitz,J. C., Casstevens,T. M., Lu,F., Harriman,J., Elshire,R. J. et al., 2014 TASSEL-GBS: A High Capacity Genotyping by Sequencing Analysis Pipeline. PLoS ONE 9: e90346.

Gualdrón Duarte,J. L., Bates,R. O., Ernst,C. W., Raney,N. E., Cantet,R. J. et al., 2013 Genotype imputation accuracy in a F2 pig population using high density and low density SNP panels. BMC Genet 14:38.

Heffelfinger,C., Fragoso,C. A., Moreno,M. A., Overton,J. D., Mottinger,J. P. et al., 2014 Flexible and scalable genotyping-by-sequencing strategies for population studies. BMC Genomics 15: 979.

He,J., Zhao,X., Laroche,A., Lu,Z.-X., Liu,H. et al., 2014 Genotyping-by-sequencing (GBS), an ultimate marker-assisted selection (MAS) tool to accelerate plant breeding. Front Plant Sci 5: 484.

Honsdorf,N., March,T., Hecht,A., Eglinton,J., and Pillen,K., 2014 Evaluation of juvenile drought stress tolerance and genotyping by sequencing with wild barley introgression lines. Mol Breeding: 1–21.

Huang,Y.-F., Poland,J. A., Wight,C. P., Jackson,E. W., and Tinker,N. A., 2014 Using genotyping-by-sequencing (GBS) for genomic discovery in cultivated oat. PLoS ONE 9: el02448.

Hyma,K. E., Barba,P., Wang,M., Londo,J. P., Acharya,C. B. et al., 2015 Heterozygous Mapping Strategy (HetMappS) for High Resolution Genotyping-By-Sequencing Markers: A Case Study in Grapevine. PLoS ONE 10: e0134880.

Johnson,J. L., Wittgenstein,H., Mitchell,S. E., Hyma,K. E., Temnykh,S. V. et al., 2015 Genotyping-By-Sequencing (GBS) Detects Genetic Structure and Confirms Behavioral QTL in Tame and Aggressive Foxes (I/ulpes vulpes). PLoS ONE 10: e0127013.

Kawahara,Y., de la Bastide,Melissa, Hamilton,J. P., Kanamori,H., McCombie,W. R. et al., 2013 Improvement of the Oryza sativa Nipponbare reference genome using next generation sequence and optical map data. Rice 6: 4.

Kearsey,M. J., and Farquhar,A. G., 1998 QTL analysis in plants; where are we now? Heredity 80: 137142.

Krishnan S,G., Waters,Daniel L E, and Henry,R. J., 2014 Australian wild rice reveals pre-domestication origin of polymorphism deserts in rice genome. PLoS ONE 9: e98843.

Li,H., and Durbin,R., 2009 Fast and accurate short read alignment with Burrows-Wheeler transform. Bioinformatics 25: 1754–1760.

Lin,M., Cai,S., Wang,S., Liu,S., Zhang,G. et al., 2015 Genotyping-by-sequencing (GBS) identified SNP tightly linked to QTL for pre-harvest sprouting resistance. Theor Appl Genet 128: 1385–1395.

Liu,H., Bayer,M., Druka,A., Russell,J. R., Hackett,C. A. et al., 2014 An evaluation of genotyping by sequencing (GBS) to map the Breviaristatum-e (ari-e) locus in cultivated barley. BMC Genomics 15: 104.

Loman,N. J., Misra,R. V., Dallman,T. J., Constantinidou,C., Gharbia,S. E. et al., 2012 Performance comparison of benchtop high-throughput sequencing platforms. Nat. Biotechnol. 30: 434–439.

Lu,F., Lipka,A. E., Glaubitz,J., Elshire,R., Cherney,J. H. et al., 2013 Switchgrass genomic diversity, ploidy, and evolution: novel insights from a network-based SNP discovery protocol. PLoS Genet. 9:el003215.

Poland,J. A., Brown,P. J., Sorrells,M. E., and Jannink,J.-L., 2012 Development of high-density genetic maps for barley and wheat using a novel two-enzyme genotyping-by-sequencing approach. PLoS ONE 7: e32253.

Poland,J. A., and Rife,T. W., 2012 Genotyping-by-Sequencing for Plant Breeding and Genetics. Plant Genome 5: 92–102.

Pootakham,W., Jomchai,N., Ruang-Areerate,P., Shearman,J. R., Sonthirod,C. et al., 2015 Genome-wide SNP discovery and identification of QTL associated with agronomic traits in oil palm using genotyping-by-sequencing (GBS). Genomics 105: 288–295.

R Development Core Team, 2008 R: A Language and Environment for Statistical Computing, Vienna, Austria. http://www.R-project.org.

Rabbi,I. Y., Hamblin,M. T., Kumar,P. L., Gedil,M. A., Ikpan,A. S. *et al.*, 2014 High-resolution mapping of resistance to cassava mosaic geminiviruses in cassava using genotyping-by-sequencing and its implications for breeding. Virus Res. 186: 87–96.

Ramos,J. M., Furuta,T., Uehara,K., Chihiro,N., Angeles-Shim,R. B. et al., 2016 Development of chromosome segment substitution lines (CSSLs) of Oryza longistaminata A. Chev. & Rohr in the background of the elite japonica rice cultivar, Taichung 65 and their evaluation for yield traits. Euphytica: 1–13.

Rice,P., Longden,I., and Bleasby,A., 2000 EMBOSS: the European Molecular Biology Open Software Suite. Trends Genet 16: 276–277.

Romay,M. C., Millard,M. J., Glaubitz,J. C., Peiffer,J. A., Swarts,K. L. et al., 2013 Comprehensive genotyping of the USA national maize inbred seed bank. Genome Biol 14: R55.

Rowan,B. A., Patel,V., Weigel,D., and Schneeberger,K., 2015 Rapid and Inexpensive Whole-Genome Genotyping-by-Sequencing for Crossover Localization and Fine-Scale Genetic Mapping. G3 5: 385–398.

Sen,S., and Churchill,G. A., 2001 A Statistical Framework for Quantitative Trait Mapping. Genetics 159: 371–387.

Sonah,H., O′Donoughue,L., Cober,E., Rajcan,I., and Belzile,F., 2015 Identification of loci governing eight agronomic traits using a GBS-GWAS approach and validation by QTL mapping in soya bean. Plant Biotechnol J 13: 211–221.

Spindel,J., Wright,M., Chen,C., Cobb,J., Gage,J. et al., 2013 Bridging the genotyping gap: using genotyping by sequencing (GBS) to add high-density SNP markers and new value to traditional bi-parental mapping and breeding populations. Theor Appl Genet 126: 2699–2716.

Swarts,K., Li,H., Romero Navarro,J. Alberto, An,D., Romay,M. C. et al., 2014 Novel Methods to Optimize Genotypic Imputation for Low-Coverage, Next-Generation Sequence Data in Crop Plants. Plant Genome 7.

Takagi,H., Abe,A., Yoshida,K., Kosugi,S., Natsume,S. et al., 2013 QTL-seq: rapid mapping of quantitative trait loci in rice by whole genome resequencing of DNA from two bulked populations. Plant J 74:174–183.

Wang,L., Hao,L., Li,X., Hu,S., Ge,S. et al., 2009 SNP deserts of Asian cultivated rice: genomic regions under domestication. J Evol Biol 22: 751–761.

